# LOAM: A family of genomic language models trained on long-read soil metagenomes

**DOI:** 10.64898/2026.09.28.755120

**Authors:** Quentin RV. Ferry, Maurice Frank, Julia Jehn, Matthias Schaks, Bruno R. Steinkraus, Timothy Rajakumar

## Abstract

Soil ecosystems represent a vast, largely uncharacterised reservoir of microbial diversity. Metagenomic assembly has unlocked access to this resource, and recent advances in long-read sequencing have improved the recovery and quality of microbial genomes from complex samples. While genomic language models have proven highly effective at capturing biological concepts from large sequence datasets, they have predominantly been trained on reference genomes or short-read assemblies. Here, we present LOAM, a family of decoder-only genomic language models ranging from 25 to 624 million parameters and trained exclusively on Oxford Nanopore long-read environmental metagenomes. LOAM model performance scales predictably with model size and training-token budget. Despite a relatively small training sequence corpus, LOAM models out-performed comparably sized models across biological benchmarks, and achieved performance competitive with substantially larger state-of-the-art models trained on much larger datasets. Context-intervention experiments further showed that LOAM models use genomic information over several kilobases, highlighting the potential value of increased contiguity provided by long-read metagenome-assembled genomes. For probe-based benchmark tasks, we systematically evaluated representations across hidden layers and revealed that task-relevant biological information was frequently more linearly accessible from intermediate than final model layers. Finally, we observed that variation in zero-shot variant-effect prediction was strongly associated with the presence of homologous target sequences in the pre-training corpus. Together, these results establish long-read environmental metagenomes as a viable foundation for training competitive genomic language models and demonstrate the importance of both model scale and training-corpus composition for biological generalisation.

## 1 Introduction

Soil is one of the most complex ecosystems on Earth and still largely unmapped, evidenced by the minority of taxa that can be labelled to the species level in large metagenomic studies [1, 2]. These communities play a fundamental role in planetary health, driving biogeochemical cycling [3], crop resilience [4], and terrestrial carbon sequestration [5], as well as representing a rich source of natural products of value for medical [6] and industrial processes [7]. Yet their systematic characterisation has historically been constrained by the small fraction of soil microorganisms that can be readily cultivated under laboratory conditions [8].

Metagenomic sequencing has unlocked access to this enormous diversity by enabling microbial genomes to be assembled directly from environmental samples [9]. However, metagenome-assembled genomes (MAGs) generated from short-read sequencing can remain fragmented and incomplete, particularly across repetitive or complex genomic regions [2]. Long-read sequencing can substantially improve MAG recovery and contiguity, and recent advances in Oxford Nanopore Technologies (ONT) chemistry and base calling have markedly increased sequence accuracy [10].

In parallel, recent developments in machine learning have enabled the extraction of information from increasingly large, unlabelled datasets. Large language models, in particular, have proved effective at capturing underlying patterns and concepts from massive unlabelled datasets [11]. Applied to genomics, this has led to genomic language models (gLMs) including Evo [12], Evo 2 [13], Nucleotide Transformer [14], gLM2 [15], GenomeOcean [16], ProkBERT [17] and EDEN [18]. These models have largely been trained on reference genomes and fragmented short-read datasets, and it has not yet been established whether ONT long-read datasets would be similarly suitable.

Here, we present the LOAM family of gLMs ranging from 25 to 624 million parameters and trained exclusively on taxonomically dereplicated long-read ONT environmental metagenomes. Across this controlled series, we examine the effects of model scale and training-token budget, test whether the models exploit extended genomic context, and evaluate the biological information captured by their representations using gene essentiality, enzyme function and zero-shot variant-effect benchmarks. LOAM models show predictable improvements with scale, use genomic context across kilobase distances and achieve competitive performance relative to state-of-the-art gLMs trained on much larger sequence corpora. Furthermore, we introduce the systematic analysis of probe representations across the model depth, and reveal that task-relevant biological information is frequently more linearly accessible from intermediate than final layers. Finally, we show that zero-shot variant effect performance is strongly associated with the abundance of homologous target sequences within the pre-training corpus. Together, these results establish long-read environmental metagenomes as a viable data source for gLM modelling and highlight both model scale and training-corpus composition as determinants of biological generalisation.

## 2 Results

### 2.1 Pre-training dataset characterisation

We constructed the LOAM pre-training corpus from the Micro Flora Danica long-read (MFD-LR) dataset [2]. MFD-LR comprises 14.4 Tbp of ONT R10.4.1 sequencing data collected from soil, sediment and water across 15 habitats in Denmark and assembled into 295.7 Gbp of contiguous sequence. Given the redundant representation of abundant and closely related taxa and the benefits of deduplication demonstrated in LLMs [19], we used the 15,640 species-level representative MAGs that had been constructed by dereplicating all medium and high-quality MAGs at 95% average nucleotide identity. In total, this provided approximately 67.5 Gbp (**Table 1**) of exclusively ONT-derived sequence for model pre-training. Notably, this corpus was substantially smaller than OpenGenome2, which comprises 8.84 Tbp of sequence in total, including 854 Gbp of non-redundant metagenomic sequence used to train Evo 2 [13].

**Table 1.** DNA Corpus. Partitioning of the dereplicated MFD-LR corpus after retaining contigs of at least 3,000 bp. Entire metagenome-assembled genomes (MAGs) were assigned to training, validation or test sets. Counts, sequence totals, corpus shares and storage-shard counts are reported for each split.

| split | MAG files | records | size (bp) | share of corpus (%) | shards |
| --- | --- | --- | --- | --- | --- |
| train | 14,098 | 431,081 | 60,979,593,726 | 90.29 | 421 |
| val | 755 | 27,164 | 3,258,982,651 | 4.83 | 27 |
| test | 787 | 24,458 | 3,297,023,648 | 4.88 | 24 |
| total | 15,640 | 482,703 | 67,535,600,025 | 100.00 | 472 |

A prominent feature of MFD-LR relative to other gLM pre-training corpora was its increased sequence contiguity. MFD-LR dereplicated MAG contigs had a median length of 47.7 kb and mean length of 139.9 kb, with individual contigs extending to over 1 Mbp (**Fig. 1A**). By comparison, sequences in the metagenomic component of OpenGenome2 had a median length of 2.2 kb and mean length of 4.0 kb (**Fig. 1A**). The median MFD-LR contig therefore exceeded the 8,192-nucleotide context length used to train LOAM, increasing the availability of complete context windows from contiguous genomic sequence.

**Figure 1.**
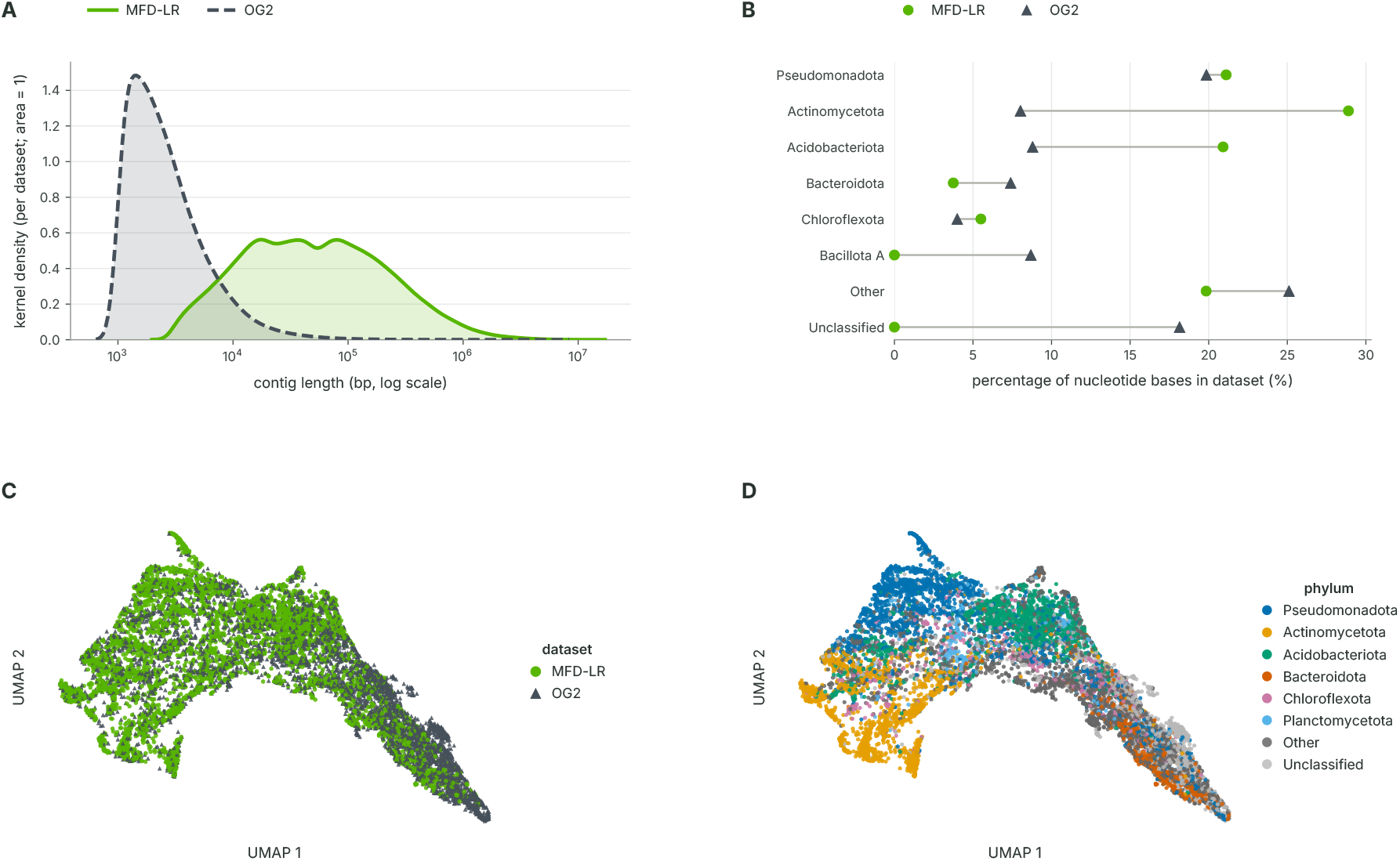
Characterisation of the LOAM pre-training corpus. **A**, Kernel density estimate of contig lengths in MFD-LR and the metagenomic component of OpenGenome2. **B,** Phylum-level composition of the two datasets. **C,D,** UMAP of 5,000 randomly sampled contigs from each dataset based on concatenated k-mer frequencies, coloured by dataset (**C**) or phylum-level taxonomic assignment (**D**).

We next compared the taxonomic composition of the MFD-LR corpus with the metagenomic component of OpenGenome2. MFD-LR was sampled from a comparatively restricted geographic region, whereas the OpenGenome2 metagenomic corpus incorporates sequences from diverse environmental and host-associated microbiomes collected globally. Despite this difference, there was considerable overlap in phylum-level representation between the two datasets although their relative abundance profiles differed (**Fig. 1B**). In particular, *Actinomycetota* and *Acidobacteriota* were more highly represented in MFD-LR whereas *Bacteroidota* and *Bacillota A* were more highly represented in OpenGenome2. To compare broader sequence compositions, we embedded a subset of contigs based on their k-mer frequency profiles (**Fig. 1C**). While both datasets showed considerable overlap, phylum-level taxonomy was associated with distinct regions of the embedding (**Fig. 1D**). Together, these analyses define MFD-LR as a substantially smaller, species-dereplicated, and more contiguous pre-training corpus that nevertheless overlaps considerably with OpenGenome2 in both phylum representation and broader sequence composition.

### 2.2 Model architecture and pre-training

We adopted a decoder-only transformer architecture based on Llama 2 [20], with full multi-head causal attention and single nucleotide tokenisation, for the LOAM family of models. During preliminary optimisation experiments, we observed a failure mode in deeper models characterised by abrupt residual-stream growth in the middle of the network. As the magnitude of the residual stream increased, subsequent block updates became insufficient to alter the residual state at training precision, effectively rendering all downstream blocks ineffective (**Sup. Fig. S1A, B**).

Importantly, this behaviour was not apparent from model loss or standard training telemetry. Similar residual stream growth has previously been described in LLMs [21], and we observed the same phenomenon upon inspection of the released Evo 1.5 7B checkpoint (**Sup. Fig. S1A, B**). To mitigate this failure mode, the final LOAM architecture incorporates peri-layer normalisation [22], together with query–key normalisation, weight decay, gradient clipping and fp32 gradient reduction. We additionally implemented targeted layer-health monitoring throughout training to proactively detect the emergence of ineffective blocks and safeguard against wasting compute on inert layers during training and at inference time.

Using this optimised architecture, we trained a family of four LOAM models spanning 25 (LOAM-25M) to 624 million (LOAM-624M) parameters (**Table 2**). Model width and depth were scaled together while maintaining an aspect ratio between 100 and 128, consistent with several widely used open-weight decoder-only LLM architectures [23, 24]. All models were trained using a causal next-token prediction objective over an 8,192-nucleotide context window. This provided a practical compromise between capturing gene-level sequence features, regulatory context, and multi-gene functional organisation such as operons and local genomic neighbourhoods, while keeping the quadratic computational cost of full self-attention tractable. The corpus splits (90.29% training, 4.83% validation and 4.88% test, see **Table 1**), global batch size and ordering, as well as optimiser schedule were held constant across models to enable controlled comparison.

**Table 2.** LOAM model series. Architecture and parameter counts for the four LOAM models. Total and non-embedding parameters are reported in millions (M); *d*_model_ denotes model width, *L* the number of transformer blocks and *d*_ffn_ the feed-forward width. All models use a head dimension of 64.

| Model | Params (M) | Non-emb. (M) | Width $d_{\text{model}}$ | Depth $L$ | Aspect $d/L$ | Heads | Head dim | $d_{\text{ffn}}$ |
| --- | --- | --- | --- | --- | --- | --- | --- | --- |
| LOAM-25M | 25.5 | 25.4 | 640 | 5 | 128.0 | 10 | 64 | 1,792 |
| LOAM-100M | 102.9 | 102.8 | 1,024 | 8 | 128.0 | 16 | 64 | 2,816 |
| LOAM-340M | 340.0 | 339.8 | 1,536 | 12 | 128.0 | 24 | 64 | 4,096 |
| LOAM-624M | 624.2 | 624.0 | 1,792 | 16 | 112.0 | 28 | 64 | 4,864 |

Every model completed a single pass through the 62.4 Gbp training corpus, with stable training loss trajectories and no abnormalities detected in layer-health monitoring (**Fig. 2A, Sup. Fig. S1C**). Final held-out validation loss after one epoch decreased monotonically with increasing model size, indicating improved modelling of the underlying sequence structure with increasing model capacity (**Fig. 2B, Sup. Fig. S2A**). To examine how model capacity interacted with training-token budget, we additionally evaluated checkpoints saved after one-third, two-thirds and the full training budget for each model. Increasing the number of training tokens consistently improved validation performance of every model in the LOAM series (**Fig. 2B, Sup. Fig. S2B**). Notably, the magnitude of this improvement increased with model capacity, with larger models achieving greater reductions in validation loss from additional training tokens than smaller models. Together, these results show that model and data scaling provide complementary benefits, with larger models not only achieving lower validation loss but also benefiting more strongly from additional training sequences.

**Figure 2.**
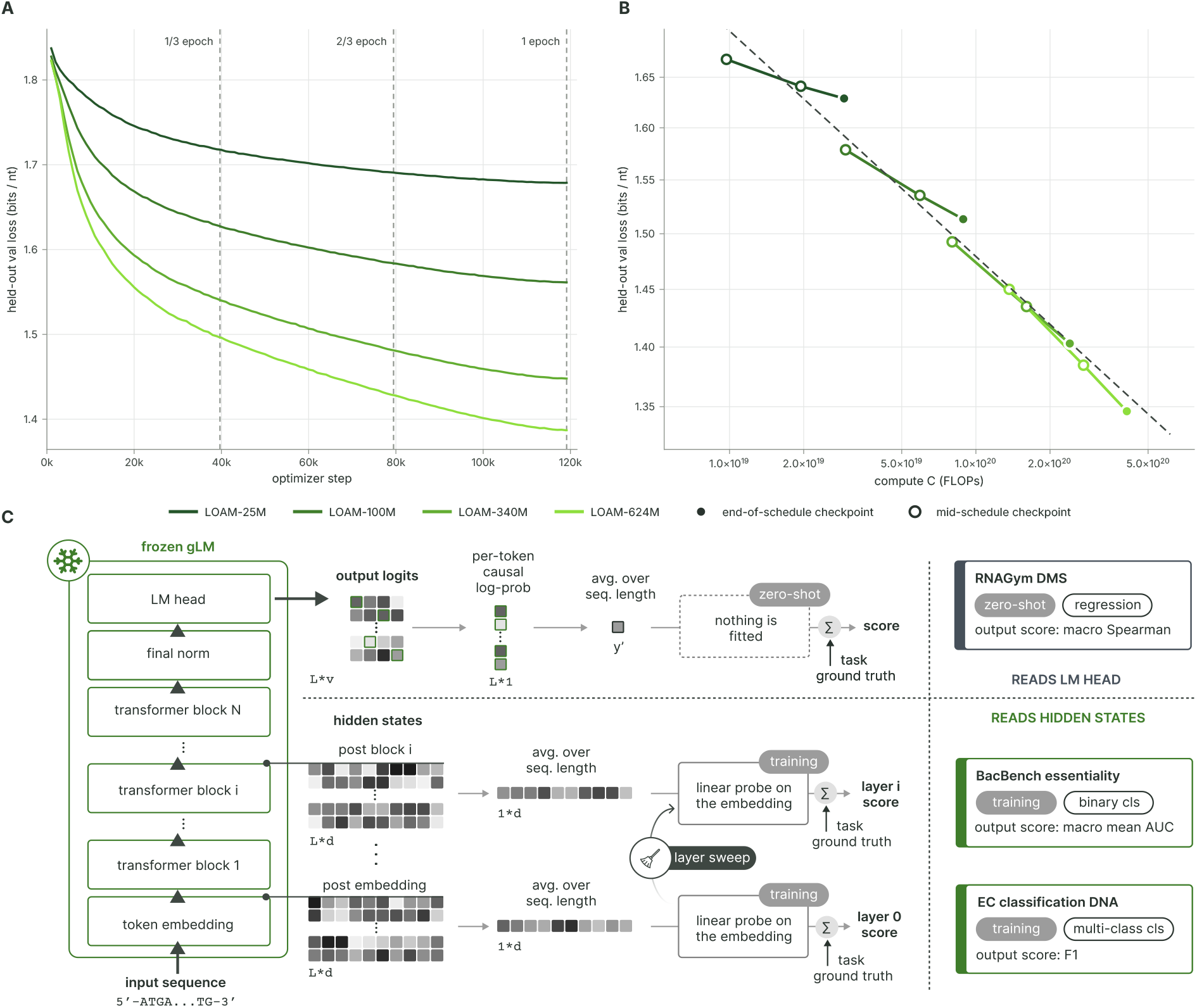
LOAM training dynamics and evaluation framework. **A**, In-loop validation loss over one epoch of training for the four LOAM models. Vertical dashed lines mark the one-third, two-thirds and full-epoch checkpoints. **B,** Held-out validation loss as a function of estimated training compute across model sizes and token budgets. The dashed line shows the fitted scaling relationship. **C,** Evaluation framework for the three biological benchmark tasks. BacBench gene essentiality and DGEB enzyme-function classification use mean-pooled hidden-state representations with linear probes and systematic layer sweeps, whereas RNAGym DMS uses model likelihoods for zero-shot variant-effect prediction.

### 2.3 LOAM models utilise extended genomic context

A potential advantage of training gLMs on long-read sequencing data is the availability of extended genomic context. We therefore tested whether LOAM models could use long-range upstream sequence information to improve prediction of downstream nucleotides. Using a set of 4 kb target sequences, we compared three conditions in which each target was provided to the model either without additional context, with 4 kb of its native upstream sequence, or with 4 kb of mismatched upstream sequence drawn from a different MAG. Loss was measured over the final 1 kb of each target sequence to quantify the effect of the upstream prefix on downstream predictions (Methods, **Fig. 3A**).

**Figure 3.**
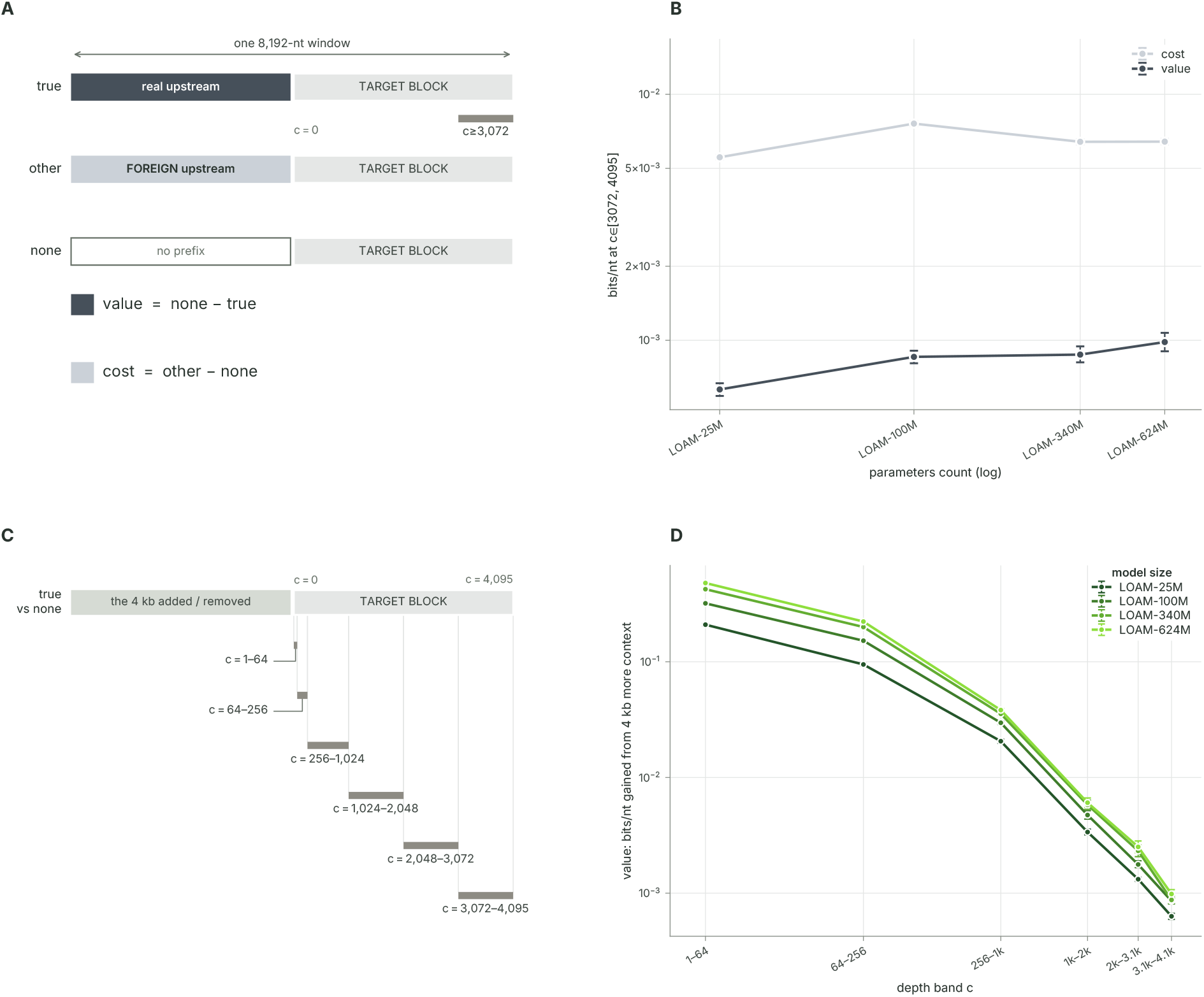
LOAM models use distal genomic context. **A**, Prefix-swap intervention comparing each target sequence preceded by its native upstream sequence, sequence from another MAG, or no prefix. Native-context value and mismatched-context cost are defined relative to the no-prefix condition. **B,** Mean value of native context and cost of mismatched context across LOAM model sizes; error bars show 95% bootstrap intervals. **C,** Target-sequence depth bands used to quantify context effects at increasing genomic distances. **D,** Predictive value of native upstream context across target depth for each LOAM model. Contextual benefit declines with distance but remains detectable over several kilobases.

Providing the native upstream sequence reduced prediction loss relative to truncation, with larger models deriving a greater benefit from this additional context (**Fig. 3B**). By contrast, replacing the native prefix with sequence from another species increased loss, and this penalty similarly grew with model size. Notably, the cost of prepending an incorrect context substantially exceeded the benefit of correct context. Because the mismatched prefix was drawn from a different species, this difference likely reflects both disruption of locus-specific dependencies and broader species-level compositional mismatch.

Having established that upstream context influenced downstream prediction, we next examined how this benefit changed over distance. We measured the reduction in loss conferred by 4 kb of native upstream context for probe regions positioned at increasing distances from the context boundary within the downstream 4 kb sequence (**Fig. 3C, D**). The benefit of native context declined progressively with increasing distance, indicating that proximal context provided the strongest predictive signal. Nevertheless, a contextual benefit remained at distances of several kilobases. In addition, larger models generally derived greater benefit from context across the range of distances examined, indicating that increasing model capacity improved context utilisation at both shorter and longer genomic distances. Taken together, these results show that LOAM models preferentially exploit proximal genomic context but also use information extending over several kilobases to inform downstream nucleotide prediction.

### 2.4 Comparator models and benchmarking tasks

For cross-model comparison, we selected gLMs trained entirely or in large part on prokaryotic sequences and spanning a broadly comparable parameter range. These comparators included gLM2 [15], GenomeOcean [16], ProkBERT [17], and Nucleotide Transformer [14]. We additionally included the larger 7 billion parameter Evo 1.5 [25] and Evo 2 [13] models as reference points representing the current gLM state of the art.

We evaluated all models using a set of complementary tasks designed to assess different forms of biological information (**Fig. 2C**). These comprised linear probing for binary classification of gene essentiality from BacBench [26], linear probing for multiclass classification of enzyme function from DGEB [27], and zero-shot prediction of RNA variant fitness from RNAGym [28]. Because the LOAM series was trained exclusively on prokaryotic sequences, analyses were restricted to prokaryotic benchmark datasets where applicable (Methods).

### 2.5 Biological information is most accessible from intermediate model layers

Linear probing assesses information encoded within a frozen model by training a simple classifier (the probe) on its hidden-state representations. Although final-layer embeddings are commonly used for this purpose, previous studies have reported improved performance from intermediate layers, making the choice of extraction layer a potentially important component of gLM evaluation [13].

We therefore extracted residual-stream representations from every hidden layer of each LOAM model and independently trained linear probes for BacBench gene-essentiality and DGEB enzymefunction classification (**Fig. 4A, B**). Probe performance varied across model depth and frequently peaked before the final layer. Among the LOAM models, an intermediate layer outperformed the final layer in 5 of 8 model–task pairs, with the performance difference between final and optimal layer greater for enzyme-function classification than for gene essentiality and most pronounced in the deeper models. Across LOAM and all comparator models, intermediate-layer representations outperformed those from the final layer in 19 of 26 model–task pairs. This effect was particularly striking for Evo2-7B, for which task-relevant information was most accessible substantially earlier in the network (**Fig. 4A, B**). Thus, evaluating only the final hidden state can underestimate the biological information encoded by a gLM and highlights the importance of systematic layer selection when benchmarking model representations.

**Figure 4.**
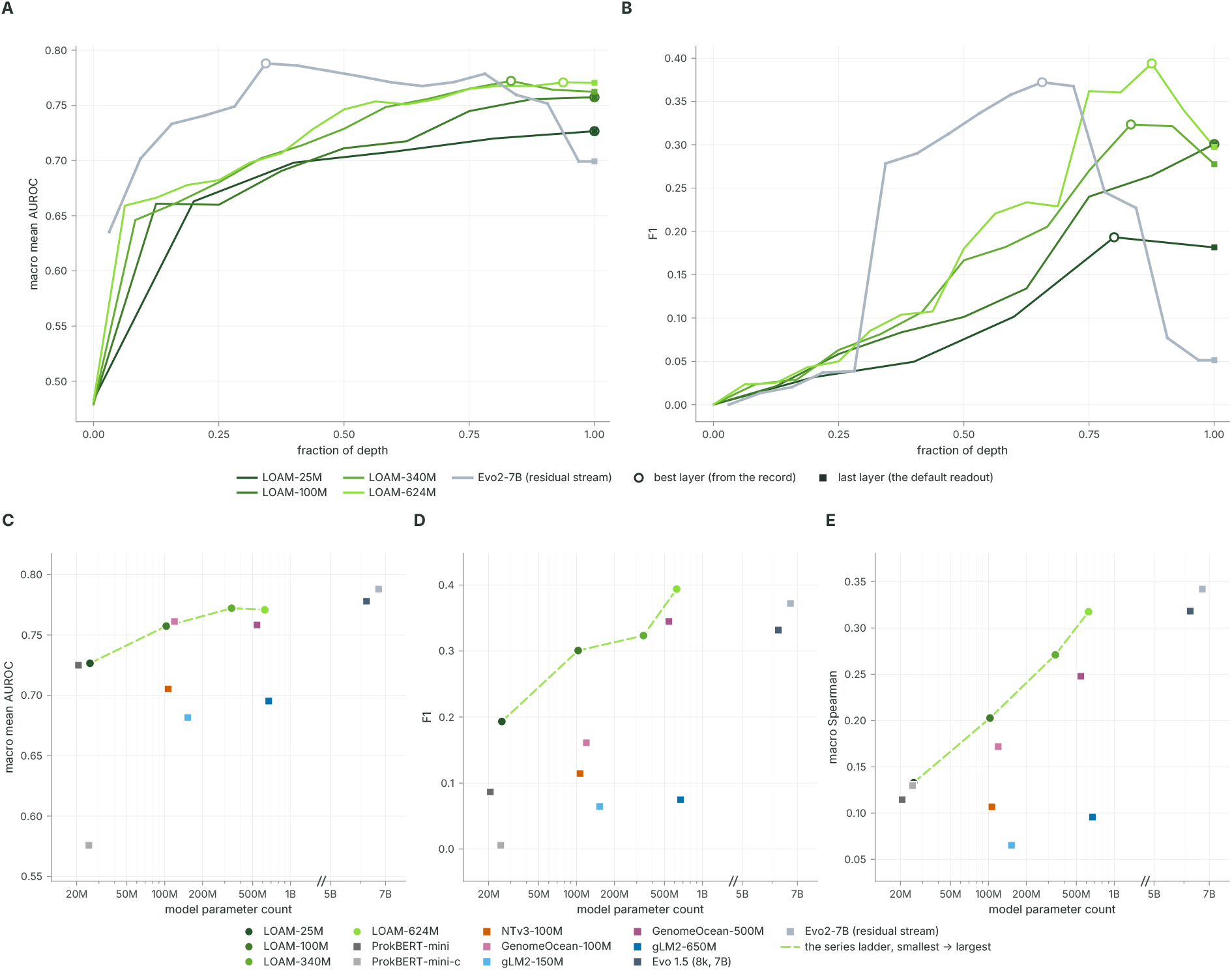
Layer-wise representations and performance across biological benchmarks. **A,B**, Linear-probe performance across normalised model depth for the LOAM series and Evo2-7B on BacBench gene-essentiality classification (**A**, macro mean AUROC) and DGEB enzyme-function classification (**B**, F1). Open circles indicate the best-performing layer and filled squares the final layer. **C–E,** Performance of LOAM and comparator gLMs as a function of model parameter count for BacBench gene essentiality (**C**), DGEB enzyme function (**D**) and RNAGym DMS zero-shot variant-effect prediction (**E**, macro Spearman correlation). For the probe-based tasks, each point reports the best-performing residual-stream layer.

### 2.6 LOAM representations enable strong prediction of gene essentiality and enzyme function

Using the best-performing hidden layer for each model, we next compared performance across gLMs. Across the LOAM series, increasing model size generally improved performance on both gene-essentiality and enzyme-function classification (**Fig. 4C, D**).

For gene-essentiality prediction, LOAM models outperformed the comparably sized ProkBERT, Nucleotide Transformer, GenomeOcean and gLM2 models across the parameter range tested (**Fig. 4C**). The larger Evo models achieved the highest overall performance, yet the competitive performance of the LOAM family indicates strong biological representation quality relative to its parameter count.

LOAM model performance was similarly strong for enzyme-function classification. Performance increased across the LOAM model series, with the largest LOAM models outperforming comparably sized gLMs, as well as exceeding the state-of-the-art Evo models (**Fig. 4D**). Together, these results demonstrate that LOAM pre-training yields intermediate representations well suited to downstream biological tasks, and that increasing model scale progressively improves the quality of these representations across both tasks.

### 2.7 Zero-shot prediction of RNA fitness

Finally, we evaluated zero-shot variant-effect prediction using the RNA fitness component of RNAGym. Because LOAM was trained exclusively on prokaryotic sequence, we restricted the analysis to 11 prokaryotic deep mutational scanning (DMS) assays spanning antibiotic resistance and gene fitness, transport, enzyme activity, and thermostability (Methods). For each variant, the frozen gLM assigned a zero-shot effect score based on the change in sequence log-likelihood relative to the corresponding wild-type sequence. We then assessed how well these model-derived scores recapitulated experimentally measured variant effects using assay-level Spearman rank correlation (**Fig. 2C**).

Zero-shot performance increased consistently with LOAM model size (**Fig. 4E**). LOAM models outperformed size-matched gLMs, achieving performance comparable to the much larger Evo 1.5 model. While Evo2-7B achieved the highest aggregate performance, LOAM-624M outperformed it on 4 of 11 individual assays (**Table 3, Sup. Fig. S3**), indicating that model scale alone did not determine assay-level performance.

**Table 3.** RNAGym DMS task breakdown. Assay-level wild-type (WT) sequence lengths, mean wild-type surprisal (bits/nt), absolute Spearman correlations (*ρ*) with measured fitness, and performance gaps (LOAM-624M minus Evo2-7B). Training-corpus coverage is reported as contigs with a detectable homologue per Gbp. The inequality denotes a lower bound where the 100,000-target search limit was reached. For BLAT_ECOLX, * denotes the Jacquier assay and ^†^ the Firnberg assay.

| assay | WT (nt) | LOAM-624M (bits/nt) | Evo2-7B (bits/nt) | LOAM-624M ($\rho$) | Evo2-7B ($\rho$) | gap | contigs w/ homologue per Gbp |
| --- | --- | --- | --- | --- | --- | --- | --- |
| IF1_ECOLI | 219 | 1.068 | 0.739 | 0.353 | 0.290 | **+0.063** | 244.1 |
| RNC_ECOLI | 681 | 1.302 | 0.597 | 0.488 | 0.412 | **+0.076** | 206.0 |
| Q837P4_ENTFA | 1,770 | 1.364 | 0.841 | 0.500 | 0.366 | **+0.134** | $\geq 1,639.9$ |
| F7YBW8_MESOW | 282 | 1.624 | 1.403 | 0.517 | 0.385 | **+0.132** | 6.5 |
| BLAT_ECOLX* | 861 | 1.631 | 0.235 | 0.416 | 0.451 | -0.035 | 24.6 |
| BLAT_ECOLX† | 861 | 1.631 | 0.235 | 0.433 | 0.439 | -0.006 | 24.6 |
| MLAC_ECOLI | 636 | 1.684 | 0.779 | 0.249 | 0.320 | -0.071 | 29.1 |
| ESTA_BACSU | 639 | 1.709 | 1.363 | 0.220 | 0.321 | -0.101 | 15.0 |
| PSAE_PICP2 | 204 | 1.892 | 1.546 | 0.125 | 0.328 | -0.203 | 0.8 |
| CCDB_ECOLI | 306 | 1.895 | 1.822 | 0.090 | 0.146 | -0.055 | 1.1 |
| BCHB_CHLTE | 156 | 1.909 | 1.541 | 0.101 | 0.306 | -0.205 | 0.0 |

We therefore investigated whether variation in relative performance was associated with representation of the assay target genes in the LOAM pre-training corpus. LOAM wild-type sequence surprisal was strongly negatively associated with performance relative to Evo2-7B (**Sup. Fig. S3A**), while the number of training contigs containing a detectable homologue of each target showed a complementary positive association (**Sup. Fig. S3B**). Accordingly, LOAM performed best relative to Evo2 on well-represented targets, whereas its largest deficits occurred for genes with few or no detectable homologues in the training corpus. Corpus representation did not, however, fully explain performance in this benchmark task. For all four assays on which LOAM-624M outperformed Evo2, Evo2 assigned the wild-type sequence higher likelihood, indicating that LOAM recovered the experimental ordering of mutations more accurately despite lower absolute familiarity with the target gene. Together, these results identify homologous sequence coverage as an important determinant of zero-shot variant-effect prediction and highlight pre-training corpus composition, alongside model scale, as a key factor in gLM biological generalisation.

## 3 Discussion

In this work, we present LOAM, a family of four gLMs trained exclusively on long-read ONT environmental metagenomic data. Using a taxonomically dereplicated corpus of MAGs derived from the Micro Flora Danica dataset, we demonstrate that ONT-derived environmental genomes are suitable for genomic language modelling. Across a controlled model-size series, LOAM models exhibited monotonic improvements in validation loss with increasing scale and captured biologically relevant information, including gene essentiality and enzyme function. These microbiome-focused models were competitive with substantially larger gLMs across prokaryotic benchmarking tasks. Zero-shot variant-effect prediction further showed that relative performance across individual targets was strongly associated with their representation within the pre-training corpus, as assessed through both wild-type sequence surprisal and homologue abundance. Together, these findings establish the feasibility of training competitive foundation models directly on long-read environmental metagenomes and highlight both model scale and training-corpus composition as important determinants of biological generalisation.

Establishing the viability of long-read sequencing derived datasets for genomic language modelling is particularly important because it unlocks the potential to use highly contiguous MAGs to train models with extended context windows. Although present models were limited to an 8,192-nucleotide context window, our context-intervention experiments demonstrate that LOAM uses sequence information across this full span. Future work could extend the effective context length through continued long-context pre-training combined with rotary position-embedding extension methods such as position interpolation or YaRN [29], as demonstrated by Evo [12] and Carbon [30]. Beyond simply increasing the context length, it will be important to develop dedicated benchmark tasks that depend on long-range interactions such as operon organisation and biosynthetic gene-cluster identification, in order to more closely evaluate gLM distal context utilisation.

Our results also emphasise the importance of training-corpus composition. Metagenomic datasets pose a particular challenge from extensive sequence redundancy arising from both evolutionary conservation and the uneven abundance of taxa within sampled communities. Duplicate or near duplicate data can reduce the effective diversity of the training corpus and bias learning towards highly represented phylogenetic branches. Published gLMs have addressed this problem using different strategies, including sequence and protein cluster-based filtering [12, 13] and representation-based semantic deduplication [15], however, the optimal method to reduce redundancy remains unclear. In this work, we instead trained on a taxonomically dereplicated MAG corpus. This represents a biologically guided means of reducing repeated representation of intra-species genomes while retaining homologous genes and conserved genomic elements across distinct species. This distinction is particularly relevant to our zero-shot results, where assay-level performance was strongly associated with the representation of homologous target sequences within the training corpus.

A broader methodological finding of this work is that biological information was frequently most accessible from intermediate rather than final hidden layers within a gLM. Across two probing tasks and multiple models, systematic layer analyses revealed substantial variation in probe performance across network depth, with the optimal representation often occurring before the final layer. This pattern is consistent with earlier layer-probing results in Nucleotide Transformer [14]. We hypothesise that this organisation reflects the progressive transformation imposed by the next-token prediction objective, with late representations becoming increasingly specialised towards prediction of the subsequent token. While this finding would support treating the extraction layer as an evaluation parameter in future gLM benchmarking, we note that our optimal layers were selected retrospectively using the test metric. Future evaluations should use independent validation sets for layer selection to most accurately estimate generalised performance for a given benchmark task.

The strong performance of LOAM on gene essentiality and functional classification also highlights a potential practical application of microbial gLMs. Rapid genome sequencing is increasingly enabling the identification of previously uncharacterised microorganisms, but translating these sequences into experimentally actionable biological hypotheses remains challenging. gLMs could complement existing annotation pipelines by prioritising gene function and essentiality directly from genomic sequence, including in organisms for which experimental annotation is lacking. In pathogenic organisms, predicted essential genes may highlight candidate biological bottlenecks for subsequent experimental validation, reducing the search space for antimicrobial target discovery.

More broadly, these potential applications illustrate the value of moving from task-specific genomic predictors towards general sequence representations that can support multiple downstream biological tasks. Large language models have demonstrated how large-scale pre-training can enable such a transition, and genomic language modelling may offer an analogous opportunity in computational biology. From this perspective, models such as LOAM are not endpoints, but general sequence representations upon which task-specific fine-tuning, functional prediction, sequence generation, and ultimately closed-loop model-to-experiment workflows can be built. Environmental long-read metagenomes are particularly attractive foundations for such models because they cover a breadth of biological diversity that remains poorly represented in current reference databases. As sequencing expands access to this diversity, and as genomic models increase in scale and effective context length, gLMs may provide an increasingly general computational interface between biological sequence and function.

## 4 Methods

### 4.1 Pre-training corpora characterisation

OpenGenome2 was downloaded from Hugging Face, and dereplicated MFD-LR MAGs were obtained from Zenodo [31]. Contig lengths were measured for the complete datasets and used to calculate kernel density distributions. For taxonomic and sequence-composition analyses, a seeded random subset of 5,000 contigs was selected from each dataset. GTDB [32] (version R220) taxonomy for MFD-LR contigs was obtained from supplied metadata. OpenGenome2 contigs were classified using MMseqs2 [33, 34] (easy-taxonomy, `--lca-mode 3, --majority 0.5, --tax-lineage 1`) using a GTDB (version R220) reference, with sequences lacking a phylum-level label retained as unclassified. Phylum composition was calculated as the percentage of nucleotide bases assigned to each phylum.

The same subset of 5,000 contigs was used to compare sequence composition by k-mer frequency. Canonical k-mer frequencies, in which each k-mer was combined with its reverse complement, were calculated for *k* = 4, 5, 6, and concatenated into a single feature vector per contig. These vectors were embedded using UMAP [35] (`umap-learn, n_neighbors=20, min_dist=0.1, metric= “euclidean”, random_state=42`).

### 4.2 Dataset segmentation

Dereplicated MFD-LR MAGs were filtered to retain contigs ≥3,000 bp, yielding 15,640 MAGs, 482,703 contigs and 67.5 Gbp of sequence. Entire MAGs were randomly assigned to training, validation or test partitions before model training, preventing sequences from the same MAG from appearing across splits. The same partitions were used to train all four LOAM models. The resulting splits contained 61.0 Gbp (90.29%) for training, 3.26 Gbp (4.83%) for validation and 3.30 Gbp (4.88%) for testing (**Table 1**).

### 4.3 Tokenisation, sequence augmentation and packing

DNA sequences were tokenised at nucleotide resolution using a 58-token vocabulary containing canonical bases, IUPAC ambiguity codes, structural tokens and reserved control tokens. Ambiguity codes were retained as distinct symbols rather than collapsed to N.

Sequences were divided into 8,192-nt windows with a stride of 7,992 nt, resulting in a 200-nt overlap between consecutive full-length windows. One trailing remainder was retained for each contig when the final window was shorter than 8,192 nt. These windows constituted the stored training records. During training, each record was reverse-complemented with probability 0.5 using the IUPAC complement table and was therefore presented once per epoch in either its stored or reverse-complemented orientation. The same augmentation was applied during in-loop validation.

Records were packed into 8,192-token sequences using a best-fit strategy with no padding. Fulllength records generally occupied a complete pack, whereas shorter trailing records were combined where possible and separated by end-of-sequence (EOS) tokens. When no buffered record fit the remaining capacity of a pack, a record could be split across packs, with the continuation treated as a new document. Attention was causal and restricted to tokens within each packed document using the FlashAttention-2 variable-length kernel [36], with rotary positions reset at document boundaries (see Section 4.4). Targets corresponding to the first token of each document and to EOS separators were excluded from the loss.

### 4.4 Model architecture

LOAM comprises four decoder-only causal transformers based on the Llama 2 decoder block [20], with root-mean-square normalisation [37], rotary position embeddings [38], multi-head causal attention, SwiGLU feed-forward networks [39], bias-free linear layers and untied input embeddings and output heads. The complete architecture consists of an embedding layer, a stack of transformer blocks, final root-mean-square normalisation and a linear output head.

Two modifications were made to the Llama 2 block. First, peri-LN [22] was used, with normalisation applied both before each attention or feed-forward branch and before its output was merged into the residual stream. Second, queries and keys were normalised using per-head root-mean-square normalisation [40] after rotary position embedding.

The four LOAM models span 25–624 million parameters (**Table 2**). Model width increased from 640 to 1,792 and depth from 5 to 16 layers, with 10–28 attention heads. Head dimension was fixed at 64 across all models, and full multi-head attention was used. Feed-forward width followed the Llama 2 convention of approximately (8/3) times the model width. Rotary position embeddings used a base of 10,000.

Linear and embedding weights were initialised from a normal distribution with standard deviation 0.02, and normalisation gains were initialised to one. Attention and feed-forward output projections were scaled by the inverse square root of twice the layer count to limit residual-stream growth with depth.

### 4.5 Regularisation

Query–key normalisation, gradient clipping at a global norm of 1.0, output-logit z-loss with coefficient 1 × 10^−4^ [41], and decoupled weight decay [42] were applied during training. The z-loss was calculated as the specified coefficient multiplied by the mean squared log-partition function of the output logits over scored positions and was excluded from reported validation and held-out losses. Weight decay was set to 0.1 for hidden parameters and 0.0 for embeddings, the output head, and normalisation gains.

### 4.6 Optimisation

All models were optimised using AdamW [42], with *β* = (0.90, 0.95) and *ε* = 1 × 10^−8^. A peak learning rate of 3 × 10^−4^ was used across all model sizes and parameter groups without width- or depth-dependent rescaling. The learning rate was linearly warmed up over 100 optimiser steps and subsequently cosine-annealed to 10% of the peak value at the final training step.

The global batch size was fixed at 524,288 tokens per optimiser step across all training runs. At a world size of eight, per-device batch sizes of 4–8 and gradient accumulation factors of 1–2 were used to accommodate model-dependent memory requirements while maintaining a constant global token count. Forward and backward passes were performed in bf16, while sharded master parameters, Adam moments, and distributed gradient reductions were maintained in fp32.

Training duration was defined by token budget, with one epoch over the training split corresponding to 119,134 optimiser steps for all model sizes. Intermediate checkpoints were retained at one-third, two-thirds and one epoch. A single run was performed for each model size using a shared random seed, which determined parameter initialisation, data-shard permutation, and per-window augmentation sampling. Consequently, all four models were trained on the same batches in the same order and orientations.

### 4.7 Training monitoring and layer health

Training loss, cross-entropy and z-loss components, pre-clipping global gradient norm, learning rate, step time, token throughput, model FLOPs utilisation, and device-memory usage were recorded at every optimiser step. No loss spikes, gradient excursions, stalls, or non-finite values were detected in the four LOAM training runs.

Layer health was monitored using the relative residual update 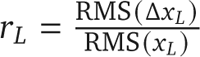 where *x_L_* and Δ*x_L_* denote the residual stream and block update at layer *L*, respectively. Values were compared with the bf16 unit roundoff, 0.003906, below which an update may not alter the represented residual state at training precision. The diagnostic was evaluated every 50 training steps using a fixed synthetic batch. Residual-stream and block-update RMS values were measured throughout the network, with the minimum *r_L_* across layers used as the primary aggregate measure. Companion diagnostics monitored optimiser second-moment statistics, weight norms, and normalisation gains. All indicators remained in a healthy range during the training of the four LOAM models.

The same residual-update analysis was performed offline on 545 held-out 16S rRNA sequences shared across models, including each reported LOAM model and two fixed pre-LN reference models (**Sup. Fig. S1**). The 16S sequences were external to the corresponding training corpora and served only as common inputs for the diagnostic. Residual states were extracted after each block, mean-pooled across tokens, and analysed in double precision to calculate residual-stream RMS, relative block updates, and cosine similarity between consecutive states. Because the offline analysis pooled representations before differencing, whereas the in-loop diagnostic calculated token-level updates before aggregation, their absolute values were not treated as interchangeable and were compared only against the common arithmetic floor (**Sup. Fig. S1**).

Validation was performed every 1,000 optimiser steps, at each predefined snapshot boundary, and at the final training step using a fixed set of 2,000 packed sequences from the validation stream. Checkpoints were saved every 1,000 steps and at snapshot boundaries corresponding to one-third, two-thirds and one epoch. Snapshot checkpoints were retained permanently, together with the final checkpoint and the checkpoint with the lowest in-loop validation loss.

### 4.8 Hardware, software and compute accounting

Model development, optimisation and training were conducted using computational resources provided by Google Compute engine and the MeluXina supercomputer at LuxProvide, Luxembourg. Training was performed on nodes containing NVIDIA A100 accelerators using fully sharded data parallelism (FSDP) [43], with parameters, gradients, and optimiser states sharded across all devices. All runs used Python 3.11.15, PyTorch 2.12.0+cu130 [44], FlashAttention 2.8.3 [36], and NCCL 2.29.7+cuda13.2.

Training the four models consumed an estimated 977.1 accelerator-hours, corresponding to 122.1 eight-GPU node-hours. Accelerator-hours were calculated from the mean steady-state optimiser-step time, excluding initial compilation and warm-up steps, multiplied by the number of optimiser steps and devices. Compute increased with model size, from 42.1 accelerator-hours for the smallest model to 521.8 accelerator-hours for the largest, with corresponding wall-clock training times of 8 and 87.5 h and mean optimiser-step times of 0.1588 and 1.9711 s, respectively.

Interrupted runs were resumed from the most recent complete checkpoint, restoring model, optimiser, scheduler, data-loader state, and random-number-generator states. Complete learning curves were retained for all model sizes (**Fig. 2A**).

### 4.9 Held-out loss measurement and scaling analysis

Two held-out loss estimates were used and analysed separately because they differed systematically in sampling. The in-loop validation loss shown in **Fig. 2A** was evaluated every 1,000 optimiser steps and at each snapshot boundary on a fixed head-of-stream subset of the validation data. Each evaluation scored 1,984 packed sequences of 8,192 tokens, corresponding to at most 16,252,928 tokens. Pure cross-entropy was reported in nats with the z-loss excluded.

All reported checkpoint losses were obtained from an offline rescore using a fixed sample shared across models and checkpoints. This sample contained 40,746 packed sequences and 333,791,232 tokens, corresponding to 10.0% of the validation split. Loss was calculated in evaluation mode using the same packing and masking scheme as training and reduced by taking the mean over all tokens. Offline rescoring was performed at three matched token budgets corresponding to one-third, two-thirds and one epoch, with identical optimiser-step boundaries across the model ladder. These checkpoints were used for the matched-budget comparisons in **Sup. Fig. S2A**. Losses were converted from nats per token to bits per nucleotide by division by ln 2.

Training compute for **Fig. 2B** was estimated as *C* = *D* · (6N + 12 · *n*_layer_ · *S* · *d*_model_), where *D* is the number of tokens consumed, *N* the non-embedding parameter count, *n*_layer_ the number of transformer blocks, *S* the sequence length, and *d*_model_ the model width. The term 6*N* accounted for the standard cost of the weight matrices, with 2*N* for the forward and 4*N* for the backward pass respectively. The second term in the parenthesis modelled the attention cost: In each layer, every token compares itself against up to *S* earlier positions (the attention scores) and then combines their value vectors (the weighted sum). Each step costs about 2 · *S* · *d*_model_ per token, so about 4 · *S* · *d*_model_ for both. Multiplying by 3 for forward plus backward and by *n*_layer_ layers gives the full term 12 · *n*_layer_ · *S* · *d*_model_.

### 4.10 Frozen-model biological evaluation

Three biological tasks (Section 4.11) were evaluated using a common harness with identical serving and evaluation logic across the LOAM series and comparator models. Each model was accessed through a registered adapter running in its native software environment and communicating tensors to a torch-free evaluation core. Adapters declared their supported capabilities and readout taps. Supported capabilities comprised embedding extraction, returning hidden states from specified layers with a specified pooling operation; sequence log-likelihood, returning a reduced causal log-probability per sequence; and masked-marginal log-likelihood ratio, calculated for each variant as the sum across mutated positions of the mutant-versus-wild-type logit difference when the corresponding wild-type position was masked [45]. Readout taps were defined either by construction or empirical probing.

Each task registered its dataset, required capability and ordered fallbacks, evaluation metric, and selection rule before model loading. Input sequences were normalised by removing whitespace, converting to uppercase, and replacing U with T. Each task then issued a single batched adapter call across its dataset, after which probe fitting and metric computation were performed on CPU. Inputs exceeding a model’s context limit were counted and handled using task-specific strategies described in Section 4.11; sequences were never silently truncated.

All pretrained model weights remained frozen. The two classification tasks fitted linear probes to pooled hidden states, whereas the zero-shot variant-effect task involved no fitted downstream model. Embedding tasks used stored hidden states, whereas likelihood-based tasks used the language-model head.

#### 4.10.1 Model serving

The LOAM models were served using the native evaluation adapter from the checkpoint produced by each training run. A beginning-of-sequence token was prepended to each input and removed before pooling or scoring, yielding a usable context of 8,191 nt from the 8,192-token training window. Inference was performed in bf16 using right-padded, length-bucketed batches containing at most eight sequences and 16,384 tokens per batch. For embedding extraction, outputs from all transformer blocks were collected in a single forward pass without applying the final model normalisation, which strongly attenuated informative high-magnitude dimensions in the residual stream. Likelihood-based readouts used the model’s native forward pass and therefore retained this normalisation.

Evo2-7B (arcinstitute/evo2_7b_base, commit hash starting in 074097e; 6,483,156,736 parameters; 8,192-nt context; bf16; single-nucleotide tokeniser) was served through the published package. Embeddings were extracted from the residual stream after 17 of the 32 blocks, comprising every second block and the final block. Variant-effect prediction used causal sequence log-likelihood with an end-of-document token prepended as context.

Evo 1.5 (evo-design/evo-1.5-8k-base, commit hash starting in 99a9a4d; 6,452,781,056 parameters; 8,192-nt context; single-nucleotide tokeniser) [25] did not expose hidden states directly; residual-stream states were therefore captured using forward hooks after each block, yielding 33 taps without model-level normalisation. Variant-effect prediction used causal sequence log-likelihood with a prepended end-of-document token.

`gLM2-650M` (`tattabio/gLM2_650M`, commit hash starting in 08754cb; 670,613,760 parameters) and gLM2-150M (`tattabio/gLM2_150M`, commit hash starting in 93c8852; 152,457,600 parameters) were evaluated with a 4,095-nt context in bf16. Their mixed nucleotide/amino-acid vocabulary is case-sensitive; DNA sequences were converted to lowercase and prefixed with the forward-strand marker, which was excluded from pooling. Hidden states were read from the model output tuple without the closing normalisation contained in the discarded model head. Layer sweeps comprised one tap per block, yielding 33 and 30 taps, respectively. Variant-effect prediction used masked-marginal log-likelihood ratios.

GenomeOcean-500M (`pGenomeOcean/GenomeOcean-500M`, commit hash starting in 473f8de; 541,109,760 parameters) and GenomeOcean-100M (`pGenomeOcean/GenomeOcean-100M`, commit hash starting in 7e55853; 119,556,864 parameters) were evaluated with a 5,000-nt context in bf16. Both checkpoints were loaded as standard Mistral decoders because the repositories contained Mistral-compatible weights while the bundled modelling code targeted an older framework version. Their byte-pair encoding produced a variable number of nucleotides per token. Sequences were wrapped in the published classifier and separator tokens, both of which were retained in mean pooling to match the published procedure. The final transformer block was hooked directly because the returned final hidden state was normalised. Variant-effect prediction used causal sequence log-likelihood, with the classifier token providing initial causal context and the separator token excluded from the loss.

ProkBERT-mini (neuralbioinfo/prokbert-mini, commit hash starting in feb2520; 20,642,181 parameters) and ProkBERT-mini-c (neuralbioinfo/prokbert-mini-c, commit hash starting in 417ebe7; 24,975,380 parameters) were evaluated with a 1,020-nt context in bf16. Both models used overlapping 6-mer tokenisation, with classifier and separator tokens excluded from pooling. Because the pre-normalisation encoder applies a final normalisation outside the transformer blocks, the final block was hooked directly. Variant-effect prediction used masked-marginal log-likelihood ratios.

NTv3-100M (`InstaDeepAI/NTv3_100M_pre`, commit hash starting in 5c685dc; 106,463,419 parameters) [46] was evaluated with a 12,288-nt context and single-nucleotide tokenisation. It was the only model served in fp32 because its blocks explicitly fix their compute datatype. The U-Net architecture operates without an attention mask and requires sequence lengths to be multiples of the downsampling factor; sequences were therefore padded individually to the next valid multiple and only equal-length sequences were batched together. The layer sweep comprised 20 taps spanning six blocks and the down-and up-sampling stages, yielding mixed-resolution states that are not directly equivalent to those of a conventional residual stack. Hidden states were taken after the skip-connection addition. Variant-effect prediction used masked-marginal log-likelihood ratios.

Across the nine external models, four used causal sequence log-likelihood for variant-effect prediction (Evo2-7B, Evo 1.5, GenomeOcean-500M and GenomeOcean-100M), similarly to LOAM, and five used masked-marginal log-likelihood ratios (gLM2-650M, gLM2-150M, NTv3-100M, ProkBERT-mini and ProkBERT-mini-c). Comparator context limits ranged from 1,020 to 12,288 nt, compared with 8,191 nt for LOAM; task-specific handling of sequences exceeding these limits is described in Section 4.11.

#### 4.10.2 Layer sweep

For the two embedding-based tasks (Section 4.11), pooled hidden states were extracted from every available readout tap in a single forward pass, and an independent linear probe with identical settings was fitted at each tap. Readout taps corresponded to the residual stream after the block residual additions and before any model-level normalisation. Where available, tap zero corresponded to the embedding output before the first transformer block. The LOAM models provided 6–17 taps across the model-size ladder, including the embedding output, while comparator tap counts followed the adapter-specific definitions described above. For cross-model visualisation in **Fig. 4**, depth was expressed as the fraction of transformer blocks applied. The best-layer score was defined as the maximum test score across valid taps. Because the best-layer score was selected retrospectively using the same test metric reported for evaluation, it represents a retrospective upper bound rather than a deployable layer-selection procedure.

All LOAM and comparator models remained frozen throughout evaluation and were accessed only through registered adapters exposing hidden states or token log-probabilities; no model was fine-tuned as part of task evaluation. All four LOAM models were evaluated at their best checkpoint (lowest in-loop validation) rather than at the final scheduled training checkpoint.

### 4.11 Benchmark tasks

Three biological benchmarks were evaluated: bacterial gene essentiality [26], enzyme-commission classification [27], and prokaryotic deep mutational scanning [28]. The first two tasks fitted linear classifiers to frozen model embeddings, whereas the variant-effect benchmark ranked variants directly from frozen language-model likelihoods. Layer-wise evaluation yielded five registered benchmark rows: four embedding-based rows and one likelihood-based row. No task applied reverse complementation or other sequence augmentation; all models received the forward-strand sequence as extracted.

#### 4.11.1 Bacterial gene essentiality

##### Provenance and scope

Code and dataset for the BacBench essential-genes task were obtained from its public repositories (code: https://github.com/macwiatrak/Bacbench; data: https://huggingface.co/datasets/macwiatrak/bacbench-essential-genes-dna). Two departures from the published protocol were made: (1) the upstream probe (a neural classifier with a torch metric) was re-derived with a scikit-learn logistic regression (*C* = 1.0, up to 1,000 iterations, no class weighting) and scikit-learn metrics (AUROC and AUPRC) [47]; (2) The validation partition, which is used only to tune and early-stop that classifier, was ignored.

##### Data

The dataset contained 193,213 labelled genes from 55 genomes, including 21,626 essential genes, partitioned into genome- and genus-disjoint splits. The fitting set comprised 136,278 genes from 37 genomes and 16 genera, including 14,167 essential genes; the evaluation set comprised 27,666 genes from nine genomes and five genera, including 3,573 essential genes. A separate validation set of 29,269 genes from nine genomes was not used.

##### Input

Inputs consisted of each coding sequence with up to 128 nt of strand-aware upstream context, clipped at contig boundaries and extracted according to the upstream procedure. Context extended upstream in sequence coordinates for plus-strand genes and downstream for minus-strand genes, without reverse complementation. Inputs averaged 1,103 nt including the upstream context, with a maximum length of 4,373,909 nt for annotations spanning entire replicons. Sequences exceeding a model’s context were divided into context-length windows with 32-nt overlap. Windows shorter than 32 nt were discarded unless this would remove the labelled gene, and window embeddings were averaged to obtain one representation per gene.

##### Model query

All windows were embedded in a single batched model query and mean-pooled over tokens. One representation per sequence and model tap was returned.

##### Fitting and scoring

Features were standardised using the fitting set, and a logistic regression was fitted with inverse regularisation strength 1.0, a maximum of 1,000 iterations, and no class weighting. Performance was calculated independently for each evaluation genome using area under the receiver operating characteristic curve, with the primary metric defined as the macro-average across genomes.

#### 4.11.2 Enzyme-commission classification

##### Provenance and scope

Code and dataset for the DGEB enzyme-commission classification task were obtained from its public repositories (code: https://github.com/TattaBio/DGEB, https://huggingface.co/datasets/tattabio/ec_classification_dna). The published torch-free evaluator was called directly, with only an unused framework-specific seeding call removed. Relative to the published protocol, sequences exceeding model context were windowed rather than truncated, and optional feature standardisation was disabled to preserve the original evaluator.

##### Data

The dataset contained 640 coding sequences distributed across 128 enzyme-commission classes, with 512 sequences used for fitting and 128 for evaluation, corresponding to four and one sequence per class, respectively. Sequence lengths ranged from 102 to 14,739 nt, with a mean of 1,678 nt.

##### Input

Published coding sequences were used as input after standard normalisation. Sequences exceeding model context were divided into context-length windows with 32-nt overlap, and window embeddings were averaged to obtain a single sequence representation. No sequences were truncated.

##### Model query

All fitting and evaluation windows were embedded in a single batched query and mean-pooled over tokens. One representation per sequence and model tap was returned.

##### Fitting and scoring

The evaluator fitted a scikit-learn logistic regression using default regularisation, a maximum of 1,000 iterations, no feature standardisation, and no cross-validation. Performance was measured using the macro-averaged harmonic mean of precision and recall across evaluation classes.

#### 4.11.3 Prokaryotic deep mutational scanning

##### Provenance and scope

Code and dataset for the RNAGym fitness benchmark were obtained from its public repositories (code: https://github.com/MarksLab-DasLab/RNAGym; data: https://huggingface.co/datasets/Marks-lab/RNAgym). The published per-assay metric implementation was used without alterations. Assays were selected programmatically from the RNAGym reference table by requiring RNA type = mRNA-coding and assay description = PROK, restricting evaluation to prokaryotic protein-coding genes.

##### Data

The resulting benchmark comprised 11 assays across 10 genes, with two assays corresponding to the same gene, and 30,658 variants in total. Coding-sequence lengths ranged from 156 to 1,770 nt (**Table 3**).

##### Input

Each input consisted of the complete normalised variant coding sequence, without flanking context or windowing. Variants exceeding a model’s context limit were skipped and counted rather than truncated. This affected only the two shortest-context comparators (ProkBERT-mini and ProkBERT-mini-c), which each excluded 697 variants corresponding to one complete assay and therefore evaluated 29,961 of 30,658 variants.

##### Model query

For causal models, variant scores were obtained from batched sequence-loglikelihood queries returning the mean per-token log-probability of the complete sequence. Model-specific beginning-or end-of-document tokens were prepended as causal context and excluded from the mean. For masked models, masked-marginal log-likelihood ratios were used as defined above, with each wild-type sequence reconstructed by reverting the corresponding mutation. All variants within an assay were required to reconstruct the same wild-type sequence.

##### Scoring

No downstream model was fitted. For each assay, performance was measured as the absolute Spearman correlation between model scores and experimentally measured fitness values. The primary metric was the unweighted mean across assays with finite correlations. Assays omitted in full because of context-length constraints were excluded from this mean rather than assigned a score of zero.

### 4.12 Distal-context intervention

The contribution of distal sequence context was evaluated on 12,000 held-out 8,192-nt windows sampled uniformly at random from MFD-LR test split contigs under three conditions: native context, distal context removed, and distal context replaced with that from another window. Each window comprised a 4,096-nt distal prefix followed by a 4,096-nt target block.

For context replacement, donor prefixes were assigned by a seeded derangement of the same window population, such that each prefix was used exactly once and the native and replaced conditions were matched in prefix composition and position. Donor and recipient windows were required to originate from different contigs but were not constrained to different genomes or dereplication clusters. For context removal, the 4,096-nt target block was evaluated alone from position zero; this condition therefore altered both contextual information and absolute sequence position. By contrast, context replacement preserved sequence length and target positions while disrupting only the native prefix–target pairing.

Context value was defined as the increase in loss after removing the native prefix, and context cost as the additional increase after replacing the removed prefix with a foreign prefix. Losses were measured in bits per nucleotide, obtained by converting nats using ln 2, from fp32 forward passes with single-precision log-softmax. The first token of each target block was excluded from scoring. Effects were calculated across six target-distance intervals within the target block (1–64, 64–256, 256–1,024, 1,024–2,048, 2,048–3,072 and 3,072–4,095 nt), with the deepest interval used for the principal reported comparison. Uncertainty was estimated using 20,000 paired bootstrap resamples over windows and reported as central 95% percentile intervals.

### 4.13 Assay-level coverage analysis

For the variant-effect benchmark, wild-type sequence familiarity was quantified as the mean per-token surprisal of each unmutated coding sequence in bits per nucleotide, using the same adapter and prefix convention as the corresponding benchmark evaluation. Wild-type sequences were reconstructed by reverting each variant mutation and were required to be identical across all variants belonging to the same assay.

Training-corpus coverage was quantified by searching each translated wild-type protein against a nucleotide database constructed exclusively from the training split (431,081 records; 60.98 Gbp) using tblastn [48]. Searches used an E-value threshold of (1 × 10^−5^), a maximum of 100,000 target sequences per query, and default word-size, substitution-matrix and composition-based settings. Across all queries, 468,656 high-scoring pairs were identified. Each training record was counted at most once per query regardless of the number of matching alignments; counts reaching the target limit were therefore treated as lower bounds. Absence of a detectable homologue was not interpreted as evidence of true absence. Searches used the base-corpus contig segmentation of the same training sequence content, and resulting counts were interpreted as relative coverage rather than as counts in the training serialisation used for model fitting.

Across assays, between-model performance differences were related separately to wild-type surprisal and homologue abundance using Spearman rank correlation, with tied ranks assigned to assays sharing the same gene. Two-sided significance was estimated from 200,000 Monte Carlo permutations of assay labels using an add-one estimator. Per-assay uncertainty shown for between-model differences combined the models’ Bonett–Wright [49] 95% correlation intervals in quadrature. Because both models were evaluated on the same variants, these intervals were treated as conservative upper bounds rather than independent-error estimates.

## Data availability

LOAM was trained on the publicly available dereplicated Micro Flora Danica long-read (MFD-LR) MAG catalogue [31]. OpenGenome2 and the BacBench, DGEB and RNAGym benchmark datasets are available from their respective public repositories as described in the Methods and cited references.

## Code availability

Code used for biological benchmarking and evaluation of LOAM and comparator models is available on GitHub at https://github.com/Soilytix/gLMBench.

## Model availability

The pretrained LOAM model weights, comprising LOAM-25M, LOAM-100M, LOAM-340M and LOAM-624M, are available through Hugging Face at https://huggingface.co/collections/Soilytix/loam, together with model configurations and usage documentation.

## Acknowledgements

We acknowledge the EuroHPC Joint Undertaking for awarding project ID EHPC-AIF-2026PG01-366 access to MeluXina at LuxProvide, Luxembourg. We also acknowledge the Google for Startups Cloud Program for providing Google Cloud credits that supported computational resources used in this work.

## Supplementary material

**Figure S1.**
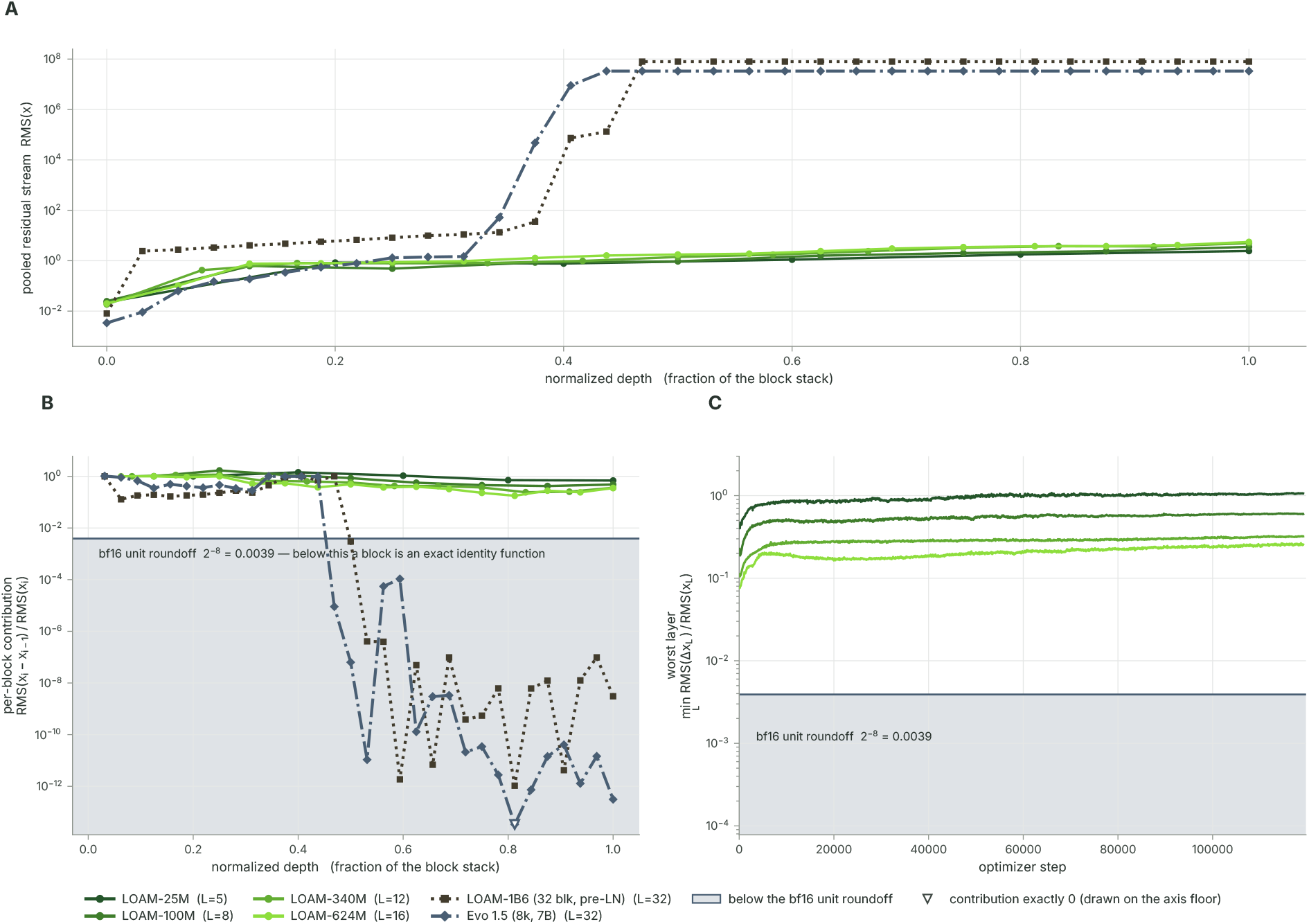
Layer collapse and health monitoring. **A**, Root-mean-square (RMS) magnitude of mean-pooled residual-stream representations across normalised model depth for the four LOAM models, compared with the preliminary LOAM-1B6 model and Evo 1.5, on 545 held-out 16S rRNA sequences. **B,** Relative residual update contributed by each block, measured as the RMS of the block update divided by the RMS of the residual state. The horizontal line marks the bf16 roundoff threshold; the shaded region indicates updates below this threshold. **C,** Minimum relative residual update across layers during LOAM training, measured every 50 optimiser steps on a fixed synthetic batch using token-level representations. All four LOAM models remain above the roundoff threshold throughout training.

**Figure S2.**
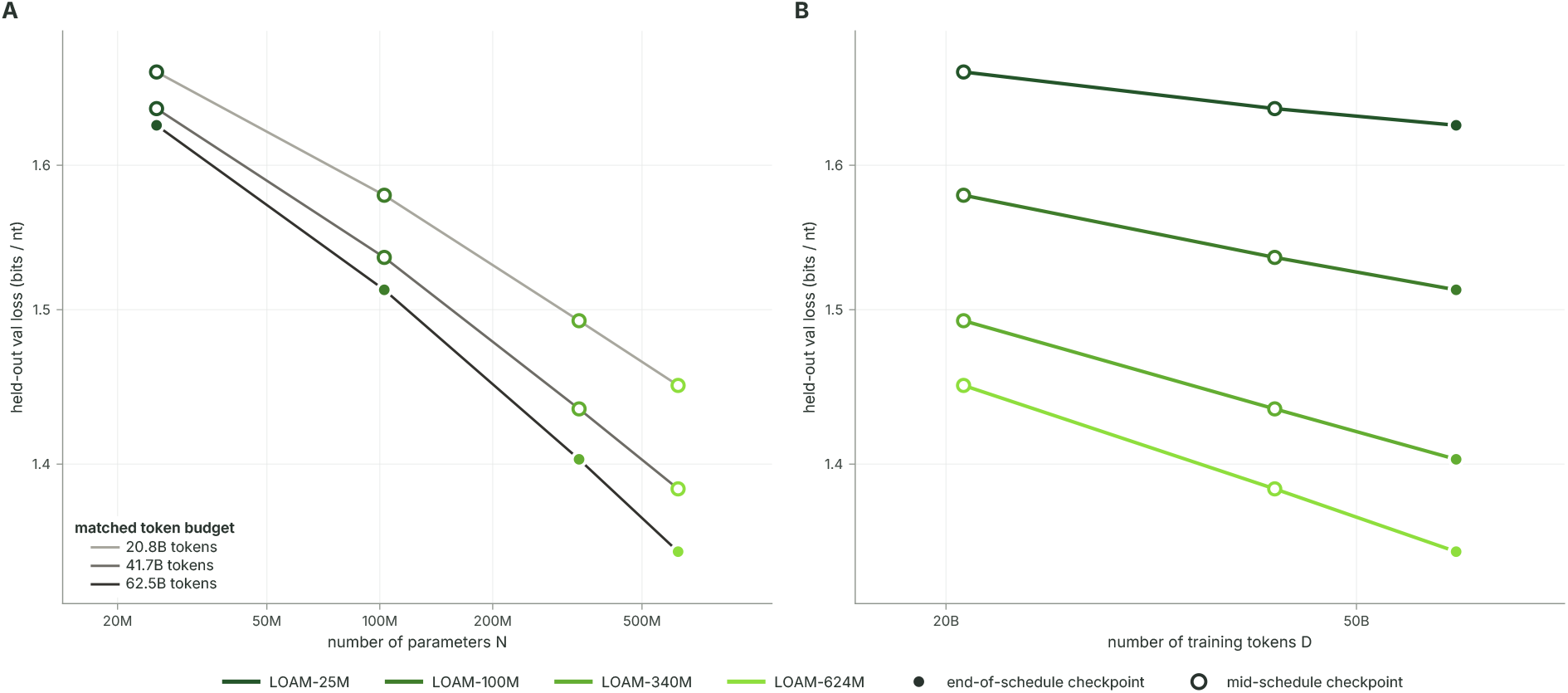
LOAM validation loss scales with model size and training tokens. **A**, Held-out validation loss as a function of non-embedding parameter count at three matched training-token budgets. **B,** Held-out validation loss as a function of training tokens for each LOAM model. Both panels use log–log axes and the same fixed validation sample across models and checkpoints. Open circles indicate intermediate checkpoints and filled circles the end of training. Larger models achieve lower loss and benefit more from additional training tokens.

**Figure S3.**
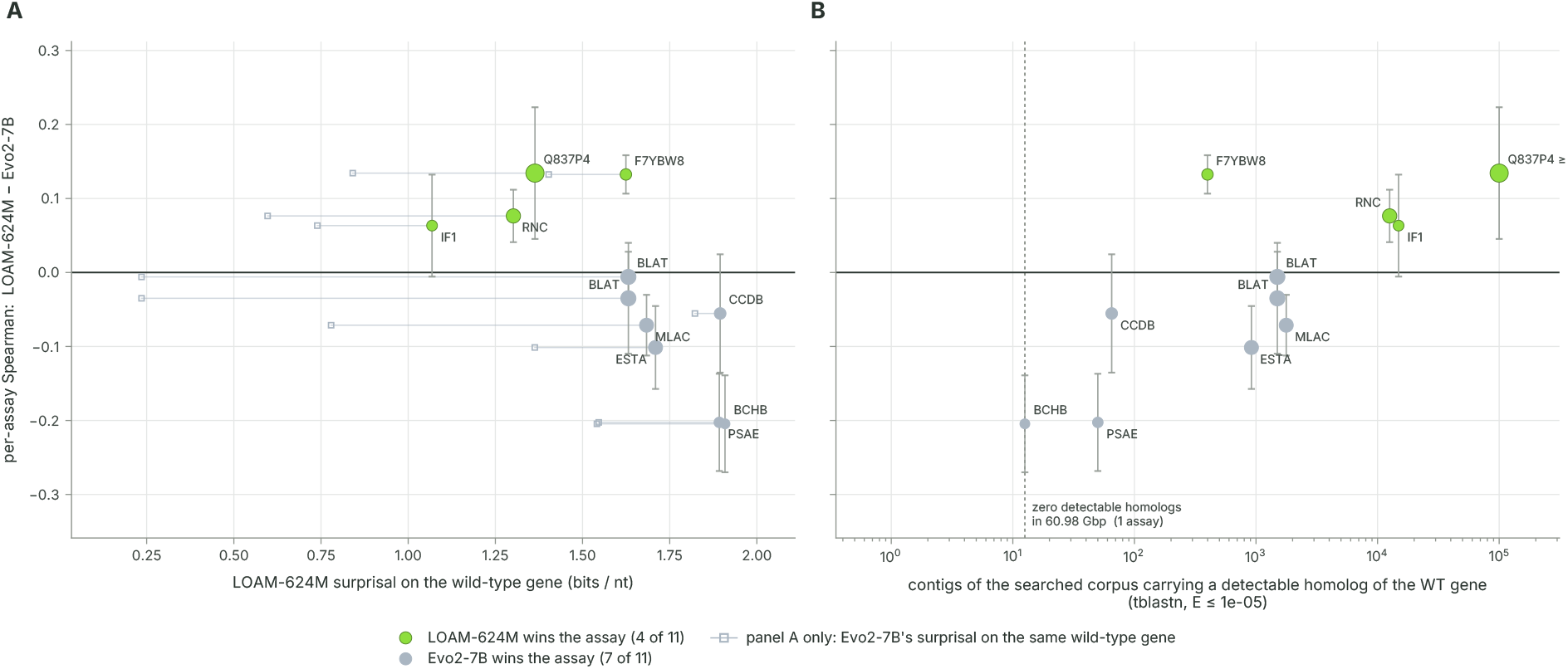
Model sequence familiarity and corpus coverage partly explain RNAGym DMS scores. **A**, Difference in per-assay RNAGym DMS Spearman correlation between LOAM-624M and Evo2-7B across 11 assays, plotted against LOAM-624M wild-type sequence surprisal. Positive values favour LOAM-624M; lower surprisal indicates greater sequence familiarity. Segments connect each marker to Evo2-7B’s surprisal for the same sequence. **B,** The same performance gap plotted against the number of training-corpus records containing a detectable homologue of the wild-type protein. LOAM’s relative performance improves with lower surprisal and greater homologue abundance. Marker area scales with the square root of coding-sequence length; error bars show conservative 95% intervals on the performance gap, combining the two models’ Bonett–Wright intervals.

